# A new map of South Manchurian mixed forests facilitates the estimation of their area for conservation purposes

**DOI:** 10.1101/2025.01.10.632318

**Authors:** Violetta Dzizyurova, Sergey Dudov, Tatyana Petrenko, Pavel Krestov, Mikhail Grishchenko, Kirill Korznikov

## Abstract

**Questions:** South Manchurian mixed forests (SMMFs) are among the most species-rich forest ecosystems in the world’s temperate zone. These forests are experiencing significant degradation and have largely been replaced by secondary forests or agricultural lands and anthropogenic non-forest vegetation. A precise quantitative assessment of the remaining forest area is crucial due to their high conservation value. This study aims to assess the current area of these forests and evaluate changes over a historical period.

**Location:** Korean Peninsula, northeastern China, and southern Primorsky Krai in Russia.

**Methods:** We used the original SMMFs distribution dataset, including vegetation relevés, literature on the distribution of these forests, and regional small-scale maps, to create a training set for classifying satellite data. We used Landsat Analysis Ready Data (1997–2001 and 2017–2021) as the basis for mapping. Mapping accuracy and area were estimated using a random stratified sampling, and forest changes were evaluated. Original relevés and literature sources were used to verify SMMFs distribution.

**Results:** The total area of SMMFs was estimated at 2,791.5±1,496.2 km², with significant uncertainty due to limited data from North Korea. Our results indicate that 45% of SMMFs are located in North Korea, 32% in Russia, 20% in China, and 3% in South Korea. Although historical declines in SMMF area have been documented, no significant changes were detected over the past 20 years. In 2021, SMMFs are highly fragmented ecosystems, primarily associated with protected areas.

**Conclusions:** We developed a high-accuracy (0.97±0.03) map of mixed forests using remote sensing techniques. This map facilitates the identification of valuable temperate forest regions and enhances understanding of current forest distribution. Our map is essential for planning restoration and conservation efforts for these critical ecosystems.

## 1. Introduction

Over the last two decades, global climate change and increasing anthropogenic pressure drive the widespread degradation of native forest ecosystems (Potapov et al. 2017; Seidl et al. 2017; Villén-Peréz et al. 2020; Diaz & Malhi 2022). The growing integration of intact forest ecosystems into global economic activities significantly diminishes their ability to provide critical ecosystem services, such as climate regulation, biodiversity conservation, carbon sequestration, and water cycle stabilization (Díaz et al. 2019; Venter et al. 2016). The loss or degradation of intact forests, as well as their replacement by secondary forests, threatens biodiversity and reduces the resilience of ecosystems (Foley et al. 2005). Temperate forests, among the Earth’s most vital biomes, are experiencing a global decline (Potapov et al. 2017; Dreiss & Volin 2020), with over 88% of broadleaf and mixed-temperate forests facing high levels of anthropogenic pressure (Mu et al. 2022). Identifying rare and vulnerable forest communities is an essential step for modern conservation efforts such as conserving the ecosystem diversity of a region (Chytrý et al. 2019). Assessing the vulnerability of ecosystems can be based on their area and rate of decline (Keith et al. 2022).

The South Manchurian mixed forests (SMMFs), belonging to the class *Quercetea mongolicae* Song ex Krestov, Dzizyurova et Korznikov 2023, are among the most species-rich forest ecosystems in the temperate zone (Krestov et al. 2006). These forests are distributed across the Korean Peninsula, northeastern China, and the southern part of Primorsky Krai in Russia (Vasiliev & Kolesnikov 1962; Kim 1990; Kolbek et al. 2003; Černý et al. 2015). SMMF plant communities are composed of coniferous species such as *Abies holophylla* Maxim. (Manchurian fir) and *Pinus koraiensis* Sieb. et Zucc. (Korean pine) with the co-dominance of the *Quercus mongolica* Fisch. ex Ledeb. (Mongolian oak), along with a diverse array of other East-Asian broadleaved tree species. Notably, *Abies holophylla* is listed as “Near Threatened” by the International Union for Conservation of Nature (IUCN) (Katsuki et al. 2013). Additionally, these forests provide crucial habitats for rare species, including the endangered Amur tiger (*Panthera tigris altaica* Temminck 1844) and the critically endangered Amur leopard (*Panthera pardus orientalis* Schlegel 1857) (Zhang & Ma 2010; Goodrich et al. 2022; Stein et al. 2024).

Unfortunately, SMMFs are undergoing significant loss and degradation, primarily due to extensive logging and fires (Dobrynin 2000). They have largely been replaced by secondary forests or anthropogenic non-forest vegetation and agricultural lands (Krestov et al. 2006). However, there are successful cases of restoration of these forests in areas previously subjected to logging and fire, provided that there are no long-term disturbances and reforestation measures are carried out (Lee et al. 2012; Kiselyova et al. 2021; Wang et al. 2021). Accurate quantification of the remaining forest area is vital, given the high conservation value of these ecosystems (Krestov & Verkholat 2003). SMMFs are reflected in the vegetation map of China (1:1,000,000) (Su et al. 2020) and the vegetation map of the Primorsky Krai (Russia) (1:500,000) (Kolesnikov 1959). Estimating the total extent and area of these forests is challenging due to their transboundary nature. Moreover, data on SMMFs in North Korea are scarce, as international access is restricted by political factors, complicating efforts to assess their distribution fully. To date, no comprehensive global estimate of SMMFs distribution or area has been conducted.

During recent decades, remote sensing and machine learning have become powerful tools to analyze global forest change. Remote sensing data are used as a standard for measuring terrestrial habitat area (Brooks et al. 2019). Mapping forests, especially mixed forests, requires detecting subtle differences in spectral images. These differences can be identified using the Phenological Metric product, developed by the Global Land Analysis and Discovery (GLAD) Laboratory at the University of Maryland. The phenological metric consists of raster files with the original values of the spectral bands, various spectral indices, and their statistics (Potapov et al. 2020a). Several regional and global geographic information products have been created based on the GLAD approach, such as the percent tree cover map of the state of Washington (Egorov et al. 2018), the global forest canopy height map (Potapov et al. 2020b), the pantropical forest structure map (Potapov et al. 2021), the global 2000-2020 land cover and land use change dataset (Potapov et al. 2022) and others.

In view of the above, this study aims to estimate the current distribution and area of SMMFs using remote sensing and to assess changes in SMMFs area.

## 2. Materials and methods

### 2.1 Study object

We define SMMFs as a regional ecosystem subgroup of the zonal mixed coniferous-broadleaved forests in the Manchurian Ecoregion (Olson et al. 2001) which reflects the warmest parts of the ecoregion (Krestov 2003; Ogureeva et al. 2012) with a monsoonal variant of the humid continental climate according to the Köppen-Geiger-Pohl system. This climate type is characterized by a pronounced summer maximum of precipitation and cold, dry winters, in which continental polar air predominates (Peel et al. 2007) (Fig. 1). SMMFs occur in northeastern China, particularly in the Changbai Mountains; on the southern slopes of the Sikhote-Alin Mountains in Russia; and in the mountainous regions of the Korean Peninsula, including the Myohyang-san, Taebaek, and Sobaek Mountains (Vasiliev & Kolesnikov 1962; Kim 1990; Kolbek et al. 2003; Černý et al. 2015) (Fig. 1). These forests are characterized by a complex assemblage of temperate tree species, with their northernmost boundary near the 44°N parallel (Krestov et al. 2006).

**Figure 1.**
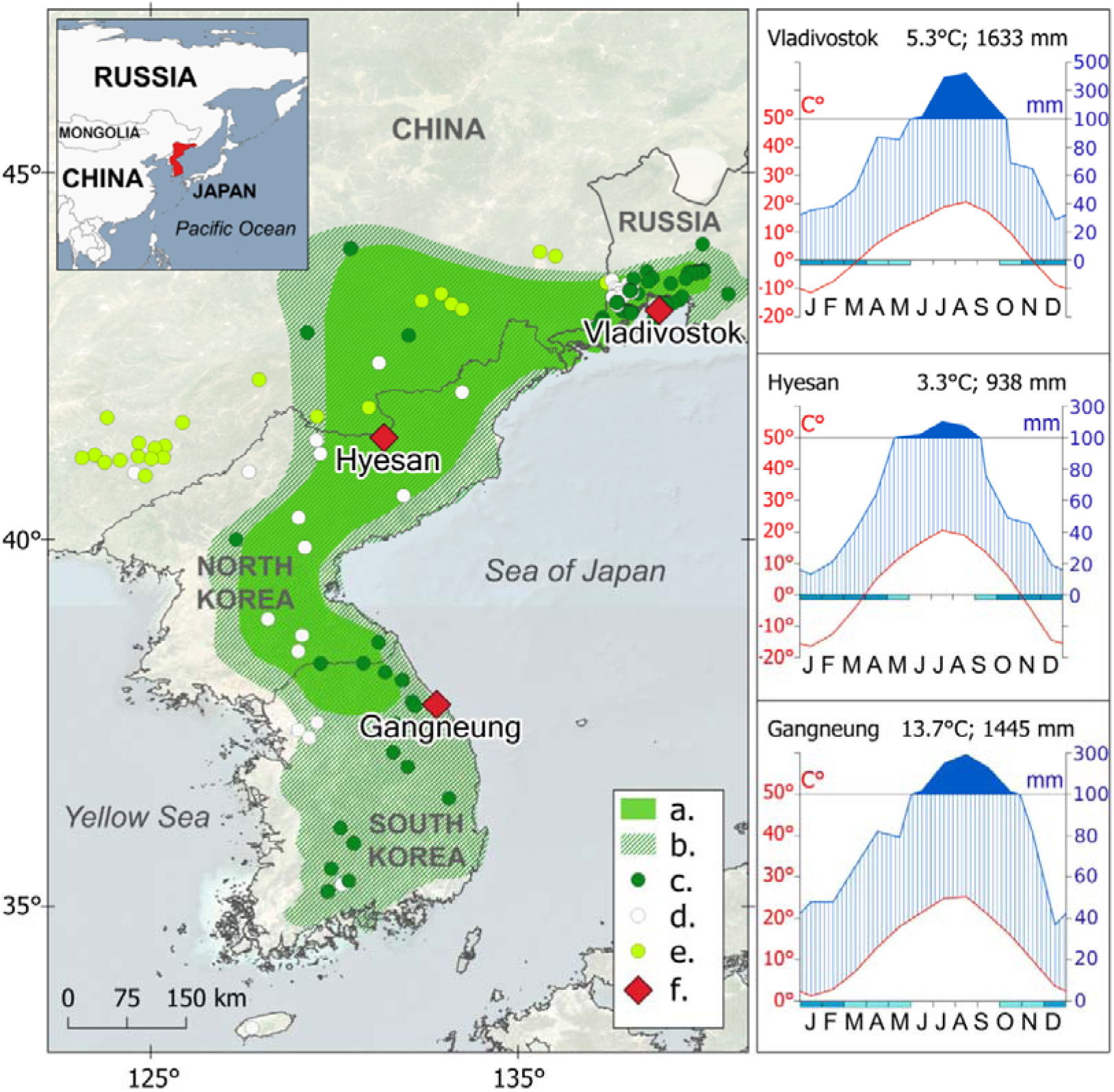
Distribution area of South Manchurian mixed forests (SMMFs) and *Abies holophylla* and climadiagramms: a – continuous distribution area of SMMFs (Vasiliev & Kolesnikov 1962); b – distribution area of *Abies holophylla* (Vasiliev & Kolesnikov 1962); c – relevés of SMMFs; d – records of *Abies holophylla*; e – presence of SMMFs according to the Chinese vegetation map (Su et al. 2020); f – wheather stations.

The forest stand is composed of ten and more tree species, comprising nemoral conifers such as *Abies holophylla* and *Pinus koraiensis* (nomenclature follows POWO, 2023), along with a variety of broadleaf species, including *Acer* spp., *Carpinus cordata*, *Juglans mandshurica*, *Kalopanax septemlobus*, *Quercus mongolica* and *Tilia amurensis*, among others. These species typically form 2–3 vertical layers (Gumarova et al. 1994; Nakamura & Krestov 2005). The shrub and herbaceous layers are well-developed (Gumarova et al. 1994; Kolbek et al. 2003), and woody lianas, such as *Actinidia* spp., *Celastrus spp., Schisandra chinensis*, and *Vitis amurensis*, are frequently encountered (Vasiliev & Kolesnikov 1962).

Like other multi-species forest ecosystems, SMMFs are characterized by natural, age-related compositional changes, which occur in cyclical patterns (Vasiliev & Kolesnikov 1962). Due to varying growth rates among coniferous and deciduous species, SMMFs undergo successional stages where deciduous species temporarily dominate the upper canopy layer during the aging process (Vasiliev & Kolesnikov 1962).

The studied forests belong exclusively to the class *Quercetea mongolicae* Song ex Krestov, Dzizyurova et Korznikov 2023, the order *Tilio amurensis–Pinetalia koraiensis* Kim ex Krestov, Dzizyurova et Korznikov 2023 and the alliance *Carpino cordatae–Abietion holophyllae* Kim ex Krestov, Dziziurova. et Korznikov 2023, as well as to the alliance *Aceri pseudosieboldiani–Quercetalia mongolicae* Song ex Takeda et al. 1994 at the southern distribution limit (Černý et al. 2015; Krestov et al. 2023).

In the southern part of their range, SMMFs are found in the upper altitudinal belts of mountain ridges and in cool, deep valleys at elevations above 1,000 m asl. In the central part, they occur across a broad range of elevations, from 0 to 700 m, while at the northern boundary, they occupy the lower altitudinal belt up to 400 m asl (Vasiliev & Kolesnikov 1962; Kim 1992; Kolbek et al. 2003; Černý et al. 2015).

### 2.2 Mapping approach

#### 2.2.1 Satellite Data

We utilized medium spatial resolution Landsat spectral data (∼30 m per pixel) provided by the GLAD laboratory. These data are available as Landsat Analysis Ready Data (ARD), which is a globally consistent 16-day time series of mosaic-normalized land surface reflectance data, covering the period from 1997 to the present. For our analysis, we selected Landsat ARD for two time periods: 1997–2001 and 2017–2021, encompassing the known SMMF range (41 1×1-degree cells, ∼368,000 km²).

We used the GLAD Phenological Metric product, which consists of annual phenological metrics derived from Landsat’s blue, green, red, near-infrared, SWIR1, and SWIR2 bands, along with their respective indices and data from a digital elevation model. Using GLAD Tools 2.0 software, we calculated annual phenological indices for the years 2001 and 2021. To address gaps in the observation series, we used data from the three preceding years to fill in any missing information. A detailed description of the GLAD methodology is provided by Potapov et al. (2020a).

#### 2.2.2 Training Data

For supervised classification, we developed a training dataset consisted of target polygons representing areas with confirmed SMMFs presence and background polygons for regions with confirmed absence. The dataset was based on SMMFs distribution data collected throughout the 21st century (the full list of sources is provided in Appendix S1). It consists of 246 phytosociological relevés and records of SMMFs presence from Primorsky Krai (Russia), northeastern China, and the Korean Peninsula, including 96 original unpublished relevés collected by the authors between 2020 and 2023 within the SMMFs range in Primorsky Krai. These vegetation relevés were gathered from 20×20-meter sample plots, capturing all vascular plant species and their cover percentages. We also used literature on SMMFs distribution and data from the Global Biodiversity Information Facility (GBIF.org, 2023) on the distribution of diagnostic, constant and dominant species.

For the forest mask, the training dataset for 2021 included 280 target polygons (∼160,000 pixels) and 315 background polygons (∼120,000 pixels). The 2001 dataset contained 300 target polygons (∼180,000 pixels) and 360 background polygons (∼330,000 pixels). For the SMMFs classification, the 2021 dataset consisted of 250 target polygons (∼30,000 pixels) and 695 background polygons (∼150,000 pixels), while the 2001 dataset included 200 target polygons (∼20,000 pixels) and 650 background polygons (∼160,000 pixels).

The operations with vector data were carried out in QGIS 3.28 (https://qgis.org/).

#### 2.2.3 Classification

We performed supervised classification of raster data using GLAD Tools 2.0 software (Potapov et al. 2020a). The classification process consisted of two consecutive steps: (1) creating a forest mask, and (2) classifying forest types (Fig. 2).

**Figure 2.**
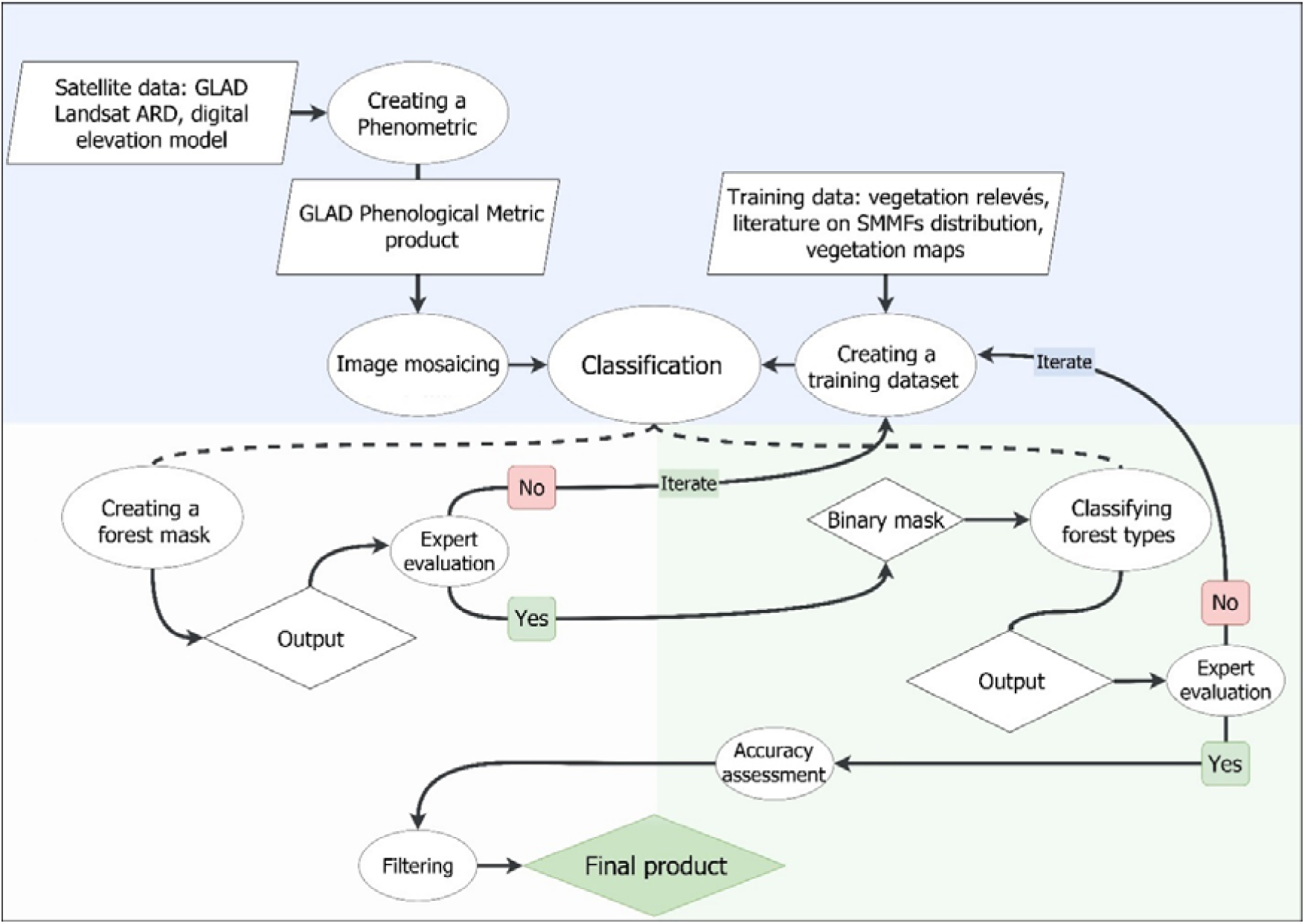
Flowchart of the processing chain for SMMFs classification using GLAD Landsat ARD.

We employed the machine learning classification algorithm provided by GLAD ARD (Potapov et al. 2020a), which is based on decision trees. A decision tree is a hierarchical classifier that determines class membership by recursively splitting the dataset into more homogeneous subsets. This splitting continues until a perfect classification tree is achieved where each terminal node contains only a single class or until predefined stopping criteria are met (Potapov et al. 2020a).

The classification output represents the calculated probability of the target class SMMFs being present in each raster cell. This was obtained using an ensemble of 25 decision trees, where each tree was built on a random sample containing 20% of the pixels from the full training dataset. The median probability of class membership was then calculated based on the outputs of all decision trees. The entire process utilized 6 parallel processes for classification, while 15 parallel processes were dedicated to building the decision tree model.

To binarize the classification results, we applied a cutoff threshold of p > 0.5, as default practice in the GLAD ARD approach (Potapov et al. 2019; 2021). The final raster product was intersected with a 10×10 km vector grid, consistent with the IUCN Red List of Ecosystems methodology (Murray 2017). We used the *exact_extract*() function from the *exactextractr* package (Baston et al. 2021) in the R programming environment (R Core Team 2023, version 4.3.1) to calculate the SMMFs area within each vector grid cell.

To evaluate mapping accuracy and track changes in forest areas over 20 years, we categorized the mapping outputs into five classes: “loss” – SMMFs were present in 2001 but absent in 2021; “gain” – SMMFs present in 2021 but absent in 2001; “stable” – SMMFs present in both 2001 and 2021; “actual area” – a composite class that includes “loss”, “gain”, and “stable” categories, excluding losses identified by the Global Forest Loss map (Hansen et al. 2013); “other forests” – a composite class encompassing all other forest types within the mapping extent.

### 2.3 Accuracy assessment

We calculated overall accuracy (OA), user accuracy (UA), and producer accuracy (PA) for each SMMF class across the entire study area and separately within the borders of Russia, China, North Korea, and South Korea. These metrics were derived using a random stratified sample of analyzed pixels following the methodology of Olofsson et al. (2014). The calculations were performed using the *olofsson*() function from the *mapaccuracy* package (Costa 2024) in the R programming environment.

Sample sizes were determined based on the proportion of each SMMF class, as well as the standard deviations of user accuracy and overall accuracy, which were estimated through preliminary analyses of random sample size (Olofsson et al. 2014, equation 12). The total sample size for the accuracy assessment was 1,064 pixels. To account for the rare “loss”, “gain”, and “stable” classes, we increased the sample sizes for these categories, as recommended by Stehman (2012), to ensure a lower standard error for the UA estimates. Specifically, 200 random pixels were allocated to each of these strata across the entire study area, with 50 pixels assigned for assessment within each country. The remaining sample pixels were allocated to the “other forest” class. The “actual area” class was evaluated by combining the “loss”, “gain”, and “stable” pixels.

Reference data for each sampled pixel were collected via visual interpretation of high-resolution satellite imagery from Google Earth™ and ESRI’s World Imagery Map, accessible via the QuickMapServices module in QGIS 3.28. Additionally, we utilized field observations and existing data on SMMFs distribution to supplement the reference data and improve training accuracy.

To minimize classification noise and remove artifacts, such as small pixel clusters at the borders of deciduous forests or conifer plantations that exhibit similar spectral responses to SMMFs, we conducted a filtering process. This process involved calculating the number of isolated patches (i.e., patches of pixels not connected at edges or corners) in the “actual area” class. We used the *get_patches*() and *lsm_p_area*() functions from the *landscapemetrics* package in R (Hesselbarth et al. 2019) to analyze patch sizes. Sequential filtering was performed using a range of size thresholds (1, 2, 5, 10, 30, 100, 500, 1000, and 5000 pixels). These size thresholds were also further analyzed to determine the level of fragmentation of the SMMFs by calculating the maximum patch size in each grid cell using the *exact_extract*() function from the *exactextractr* package in R.

To remove artifacts, we determined the required filtering threshold for each country through sampling analysis using the *olofsson*() function, then adopted this threshold for the final raster product. Artifacts below the threshold were reclassified in *terra* package (Hijmans 2020) as “other forest” and the results of the classification were compared before and after filtering for both the entire study area and individual countries. Sequential removal of small-area SMMF fragments increased user accuracy by several points but reduced producer accuracy. Finally, we adjusted the SMMF area calculations based on the optimal filtering results for each country. The following filtering thresholds were applied: 1 pixel for North Korea and 2 pixels for South Korea. For Russia and China, it was decided to use the mapping results without filtering, as the removal of individual pixels did not improve mapping accuracy (Table A1 in Appendix 1). The presence of individual pixels in these territories reflects the mosaic nature and dynamic stages of SMMF, unlike on the Korean Peninsula, where the noise is caused by mapping artifacts.

### 2.4 Assessment of the SMMFs changes

To assess changes in the SMMFs’ area over the 20-year period, we calculated the areas for each SMMF class and their 95% confidence intervals. These were based on the mapping accuracy and class proportions, following the methodology of Olofsson et al. (2014). This analysis was performed in the R programming environment using the *olofsson*() function from the *mapaccuracy* package.

To further investigate the reasons behind the changes in SMMFs, we employed a sample-based approach, as recommended by Tyukavina et al. (2015; 2022) and based on Cochran’s (1977) sampling methods. Specifically, we randomly selected 300 points from each of the “loss”, “gain”, and “stable” classes and conducted a visual interpretation of high-resolution satellite images from various years via Google Earth™. This step allowed us to cross-validate our findings with historical imagery and better understand the drivers of forest change in the study area.

Additionally, we calculated the areas of intersection between our SMMFs map products, and the Global Forest Loss and Gain maps developed by Hansen et al. (2013). This allowed us to quantify areas where the forest loss or gain trends aligned with our observations.

To evaluate historical changes in the SMMFs distribution over a longer period, we digitized all available historical cartographic sources indicating SMMFs locations. We also interpreted the records from the first forestry expeditions in the Russian part of the SMMFs range during the 1860s and 1880s. Notably, we used reports by Budishchev (1867, 1883) and the forestry expedition report of Pyastuskevich (1886), which was first published by Man’ko (2016).

We georeferenced the digitized maps to the original cartographic products in QGIS 3.28 for standardization. A 10×10 km vector grid was used to harmonize and compare data across different periods and sources.

## 3. Results

### 3.1 SMMFs area and distribution

For the first time, a large-scale map of SMMFs was created (available at <u>Zenodo</u>), reflecting the total area of SMMFs as 2791.5 ± 1496.2 km² (the evaluated area ± 95% confidence interval) in 2021. However, this estimation comes with high uncertainty, particularly for North Korea, which contributes a large amount to the global SMMFs area. The distribution of the global SMMFs area is estimated as follows: North Korea – 45%, Russia – 32%, China – 20%, and South Korea – 3% (Table 1). The overall mapping accuracy for the SMMFs was 0.97 ± 0.03 (the evaluated OA ± 95% standard error) (Table 1). Russia exhibited the highest UA (0.94) and PA (0.85) due to a more robust dataset. Conversely, results for North Korea exhibited low UA = 0.49 and PA = 0.27, reflecting challenges in data collection and georeferencing uncertainty.

**Table 1.** The evaluated area ± 95% confidence interval (CI), overall accuracy (OA), user accuracy (UA), and producer accuracy (PA) ± standard errors (SE) for each country.

| | Area, km<sup>2</sup> ± CI | OA ± SE | UA ± SE | PA ± SE |
| --- | --- | --- | --- | --- |
| **Total area** | 2791.5 ± 1496.2 | 0.99 ± 0.004 | 0.67 ± 0.02 | 0.53 ± 0.1 |
| **Russia** | 1182.4 ± 427.6 | 0.98 ± 0.01 | 0.94 ± 0.01 | 0.85 ± 0.1 |
| **China** | 746.4 ± 620.1 | 0.99 ± 0.01 | 0.55 ± 0.05 | 0.51 ± 0.3 |
| **North Korea** | 1691.5 ± 1229.4 | 0.98 ± 0.01 | 0.49 ± 0.03 | 0.27 ± 0.2 |
| **South Korea** | 104.5 ± 16.8 | 0.99 ± 0.001 | 0.51 ± 0.04 | 0.99 ± 0.001 |

There is a clear trend of SMMFs being related with protected areas (PAs). Notably: 66% of Russia’s SMMFs lies within PAs, 25% of South Korea’s SMMFs is in PAs, SMMFs in China and North Korea are also found near PAs, though exact PA boundaries are unavailable (Fig. 3a).

**Figure 3.**
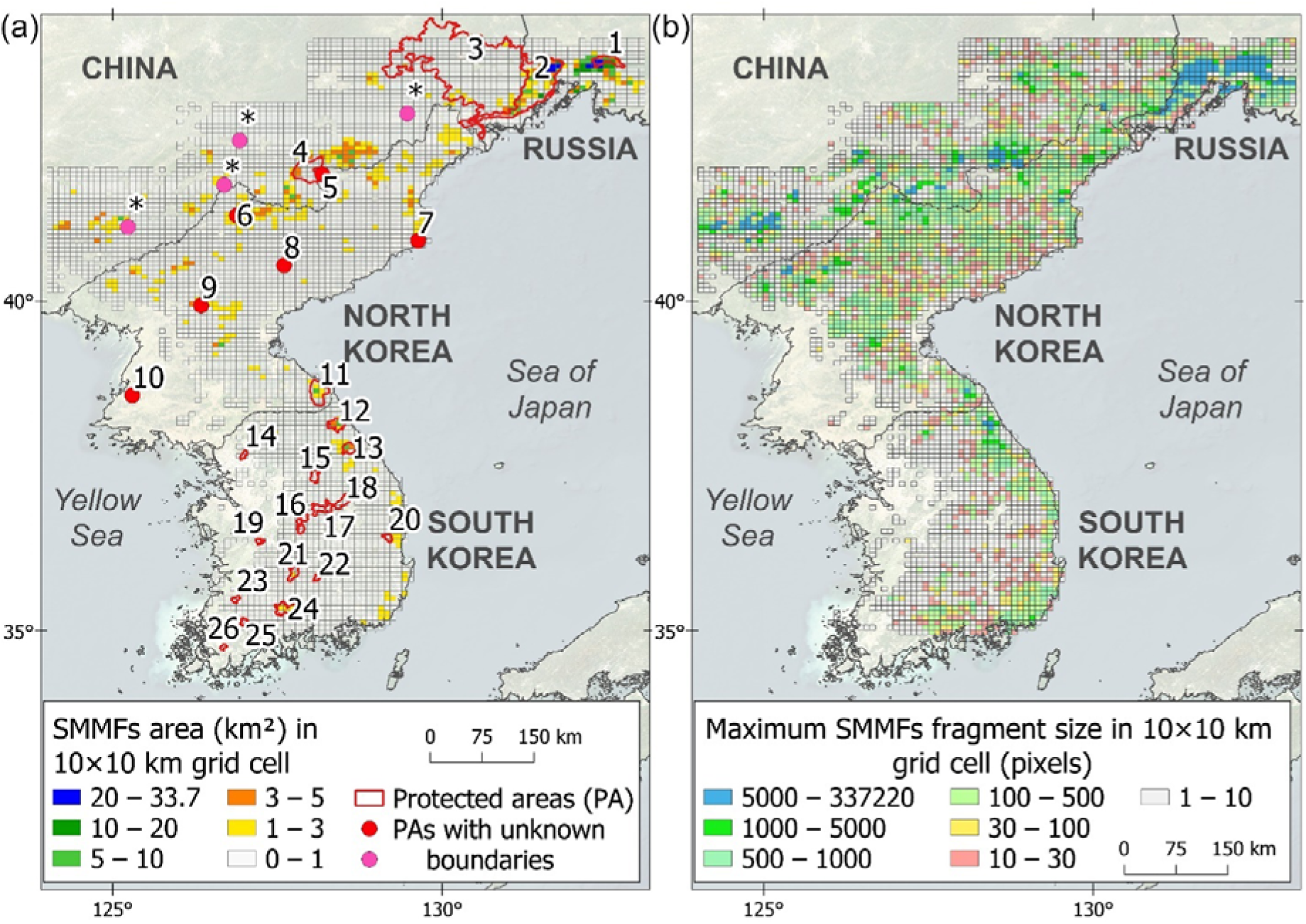
SMMFs distribution map (a) and maximum fragment value (b) in 2021 in 10×10 km grid cells. Protected areas: I. Nature Reserves: 1 – Ussuriyskiy, 4 – Changbai Mountain, 6 – Ogasan, 7 – Chilbosan, 9 – Myohyangsan. II. National Parks: 2 – Land of the Leopard National Park, including Kedrovaya Pad’ Nature Reserve, 3 – North East Tiger and Leopard National Park, 5 – Backdu-san, 8 – Lake Bujon, 10 – Guwol, 11 – Kymgan, 12 – Seoraksan, 13 – Odaesan, 14 – Bukhansan, 15 – Chiaksan, 16 – Songnisan, 17 – Woraksan, 18 – Sobaeksan, 19 – Gyeryongsan, 20 – Juwangsan, 21 – Deogyusan, 22 – Gayasan, 23 – Naejangsan, 24 – Jirisan, 25 – Mudeungsan, 26 – Wolchulsan. An asterisk (*) indicates the protected areas of China with various statuses, as listed in the database of Guo & Cui (2015).

SMMFs are highly fragmented, with larger patches (above 5000 pixels or ∼ 450 hectares) only found in Russia, primarily associated with protected areas (e.g., the Ussuriyskiy Nature Reserve and the Land of the Leopard National Park) (Appendix S2). Smaller fragments, between 1000 and 5000 pixels (90–450 hectares), are scattered across China and the Korean Peninsula, particularly in PAs like the Changbai Mountain Nature Reserve (China) and the Kumgang National Park (North Korea). In South Korea, relatively small SMMFs fragments of 100 pixels (9 hectares) or less predominate (Fig. 3b; Table 2).

**Table 2.** Average values of SMMFs fragment area for different countries.

|  | Fragment area, average value | Fragment area, median value | Proportion of 1-pixel fragments out of the total number of fragments | Proportion of 1-pixel fragments from the total area |
| --- | --- | --- | --- | --- |
| **Total area** | 4,336 pixels (390 ha) | 20 pixels (1.8 ha) | 10% | 0.002% |
| **Russia** | 52,473 pixels (4,773 ha) | 100 pixels (9 ha) | 6% | 0.0001% |
| **China** | 296 pixels (27 ha) | 20 pixels (1.8 ha) | 11% | 0.04% |
| **North Korea** | 174 pixels (16 ha) | 30 pixels (2.7 ha) | 7% | 0.04% |
| **South Korea** | 78 pixels (7 ha) | 10 pixels (0.9 ha) | 14% | 0.2% |

### 3.2 SMMF area changes in the 21st century

The area of the “loss” mapped class is 992.1 km², which accounts for 68.6% of the area of SMMFs identified for 2001. The area of the “gain” class is 833.1 km², which accounts for 72.3% of the area of SMMFs classified based on data from 2021. The class “stable area” occupies a significantly smaller area of 380.6 km² (Table A2 in Appendix 1). Interestingly, global forest loss mapped by Hansen et al. (2013) showed only 27.5 km² (2% of the area predicted for 2001) of actual SMMFs loss between 2001 and 2021, indicating that most changes in forest cover are due to classification errors rather than actual deforestation. We consider this area as a real loss of SMMFs.

Sample-based estimations of the reasons for differences in cartographic products for 2001 and 2021 yielded the following results. In the “loss” class, 17% of points are assessed as classification errors – these are dark coniferous and pine forests. 2% of points are assessed as real losses – logging for construction purposes. The remaining 81% of points, according to the results of visual interpretation of modern satellite images, are currently occupied by SMMFs. In the “gain” class, 13% are assessed as classification errors, 1% of points are restoration (increase) of SMMFs, and the remaining 86% of points are stable SMMFs. In the “stable” class, 3% of points are assessed as classification errors, and the remaining 97% of points are stable SMMFs.

## 4. Discussion

### 4.1 Map of SMMFs current distribution

The cross-border distribution of the SMMFs results in variability in the data available for training, leading to heterogeneous mapping accuracy across regions. This variability has influenced the estimation of both the current SMMFs area and its changes over the past 20 years.

Mapping mixed forests using satellite imagery is methodologically challenging due to the natural dynamics of these forests, where dominant tree species periodically change (Vasiliev & Kolesnikov 1962). As a result, even pixels from the same forest type may exhibit significantly different spectral responses, contributing to inconsistencies in the classification results for 2001 and 2021. Nguyen et al. (2023) also identified the limitations in vegetation classification of mixed deciduous-evergreen plant communities in a case study of semi-arid highlands of Morocco using a single machine learning method and suggested the need for expert validation of mapping results and ground validation.

The final estimation of the SMMFs area is a combination of mapping products from 2001 and 2021, excluding forests that have disappeared according to the Global Forest Loss Map (Hansen et al. 2013). This adjusted map aligns well with previously published sources on SMMFs distribution (Fig. 4).

**Figure 4.**
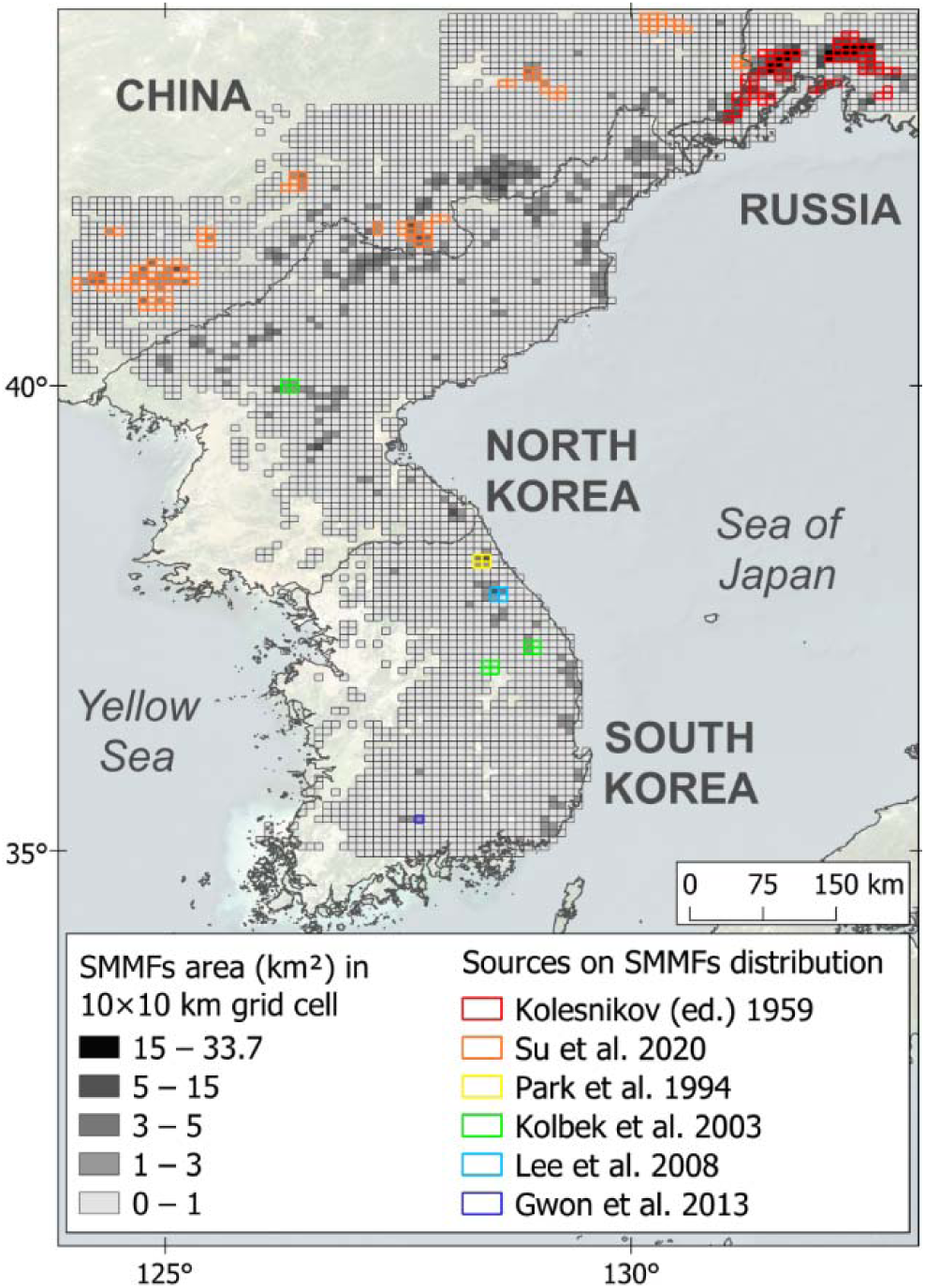
Comparison of the original mapping product with literature sources on SMMFs distribution

In Russia, SMMFs boundaries correspond with earlier literature (Vasiliev & Kolesnikov 1962; Krestov & Verkholat 2003), confirmed by numerous field observations (Appendix S1). Historically, SMMFs dominated the southern part of Primorsky Krai of Russia. Compared to China and the Korean Peninsula, the settlement and economic development of Primorsky Krai began later. This has helped preserve patches of SMMFs, particularly those currently connected with protected areas. In many cases, this relationship is the opposite: protected areas have been created, where forests have been well preserved, including those inhabited by the Amur tiger and other species of conservation concern. The largest remaining, less-fragmented SMMFs areas (above 5000 pixels or 450 hectares) are in the Ussuriyskiy Nature Reserve and its surroundings, in the Land of the Leopard National Park (on the Borisovsky Plateau and along the border with China), and in the Kedrovaya Pad’ Nature Reserve. Smaller fragments also exist around Vladivostok, but they have decreased due to logging for construction.

In China, the boundaries of SMMFs identified match well with the vegetation map of China (Su et al. 2020). The exception is a large fragment in the Yanbian Korean Autonomous Prefecture, where these forests are classified as “*Pinus koraiensis* and deciduous broadleaf mixed forest”. This classification is consistent with the presence of the southern variant of SMMF in the region. Evidence from studies (Jiang et al. 2011; Lin et al. 2024) confirms the presence of SMMF species like *Abies holophylla* and *Carpinus cordata* there. However, SMMFs in northeastern China are more fragmented than in Russia, largely due to a longer history of economic development (Yu et al. 2010; Zhao et al. 2014) and catastrophic fires (Wang et al. 2006). SMMF fragments larger than 1000 pixels (0.9 km²) are found within the Changbai Mountain Nature Reserve, in the Helong and Linjiang county cities in Jilin Province, and in the autonomous counties of Benxi Manchu and Huanren Manchu in Liaoning Province.

In North Korea, 45% of the total SMMFs area is located according to our map, which aligns with Krestov et al. (2006), who suggest widespread SMMFs presence in the country. However, precise data on SMMFs locations are scarce, except for reports of their occurrence in the Mount Myohyang area (Kolbek et al. 2003). Long-term anthropogenic pressure has significantly altered North Korea’s lowland and montane forests, fragmenting SMMFs distributions. SMMF fragments larger than 1000 pixels (0.9 km²) have been identified in the Kumgang National Park, as well as in the counties of Byokdong, P’yŏngan-bukto, and Kop’ung, Chagang-do. The country’s forest cover has drastically declined since the 1980s (Engler et al. 2014), but recent policies focus on reforestation (Lee et al. 2005; Lee & Miller-Rushing 2014). However, forming stable mixed forests in the Manchurian ecoregion can take over 1000 years (Ukhvatkina & Omelko 2016).

In South Korea SMMFs are restricted to upper altitudinal belts of mountain ranges (above 1000 m) and are more naturally fragmented than in the northern part of the range (Kolbek et al. 2003; Černý et al. 2015). Historically, SMMFs were more widespread in South Korea, but today, small fragments remain, particularly in Seoraksan National Park and other protected areas. Despite a long history of economic exploitation of the forests, these remaining fragments have high conservation value (Yu et al. 2010). Thanks to modern environmental policies, SMMFs are considered to succeed the most among the secondary communities in South Korea (Lee et al. 2012). In Odaesan National Park, SMMFs are widely distributed and are observed in other well-preserved locations in Gangwon Province, e.g., Mt. Gariwangsan, Mt. Jungwangsan, and Mt. Jeombongsan (Hong et al. 2001; Kang 2003; Lee et al. 2012).

### 4.2 SMMFs area changes in the 21st century

An evaluation of the changes in SMMFs area between 2001 and 2021 reveals minor overall differences in their total extent, although these changes exhibit strong spatial variation. The natural life cycle of trees dominants of SMMFs, particularly *Abies holophylla*, suggests that significant shifts in forest range or structure are unlikely to occur over such a short period (Vasiliev & Kolesnikov 1962; Ukhvatkina & Omelko 2016). For instance, one generation of *Abies holophylla* requires approximately 280 years from seeding to stand destruction, and the transition from the lower to the upper forest layer takes 50–60 years. This slow progression indicates that any notable differences in forest coverage are not due to dynamic processes, but rather uncertainties in the classification methods used.

These uncertainties can be attributed to the use of satellite data from different generations: the 2001 mosaic was compiled using data from Landsat 5 and 7, while the 2021 mosaic relied on data from Landsat 8. Variations in spectral resolution between these satellites influenced the accuracy of SMMFs mapping, creating discrepancies in the predicted forest areas. Despite these technological differences, an overall analysis suggests that there has been no substantial shift in SMMF distribution in the 21st century (Fig. 4). Most SMMFs loss occurred during earlier periods of forest exploitation and development (Pyastushkevich 1888; Yu et al. 2010; Kong et al. 2016).

### 4.3 SMMF area changes over the historical period

Over the last 120 years, the global area of SMMFs has significantly decreased due to extensive logging and natural disturbances such as fires (Krestov & Verkholat 2003; Yu et al. 2010). Between 1900 and 2000, human activity led to the degeneration of large tracts of SMMFs, which were subsequently replaced by secondary forests with lower biodiversity and ecosystem services (Krestov et al. 2006; Wang et al. 2021).

In Russia, major disturbances occurred between the 1860s and 1880s, when Russian settlers cleared vast areas of forests (Budishchev 1868, 1898; Pyastushkevich 1888). By 1959, the distribution of SMMFs in Russia had largely reached its current state (Kolesnikov 1959). In some regions, these forests experienced a secondary spread after earlier deforestation.

In northeastern China and on the Korean Peninsula, the SMMFs area has been drastically reduced during in the first half–mid-20th century due to extensive use associated with political events (Lee et al. 2004) when large portions of the forests were cleared (Dudin 2020). Following deforestation, secondary forests have naturally regenerated in place of SMMFs, but these secondary forests offer poorer ecosystem services and reduced productivity compared to the original SMMFs (Zhang et al. 2018).

On the Korean Peninsula, historical records trace the decline in the range of *Pinus koraiensis*, a key species of SMMFs (Kong et al. 2016). In the 1860s, *Pinus koraiensis* was distributed across the northern and eastern highlands of the Korean Peninsula, although its northern distribution was already secondary by that time. By 1930, its range in present-day South Korea had significantly diminished (Kong et al. 2016), and its distribution aligns closely with our mapping results.

South Korea harbors the southern limit of the SMMFs range. SMMFs there are restricted to the upper elevations of mountainous regions, where intact fragments persist, such as in Odaesan National Park (Lee et al. 2012). North Korea harbors the central part of the SMMF range, but widespread deforestation, particularly during the rapid urbanization period following the Korean War in 1953, has severely impacted the extent of intact forest (Lee & Miller-Rushing 2014; Lee et al. 2015).

Today, SMMFs have largely been replaced by secondary forests, such as mixed broadleaf and monodominant *Quercus mongolica* forests, which persist under the constant influence of disturbances (Dobrynin 2000; Krestov & Verkholat, 2003). Without these disturbances, oak forests have the potential to naturally revert to primary SMMFs over time (Dobrynin 2000). Restoration efforts can benefit from the adaptive capacities of key SMMFs species like *Abies holophylla* and *Pinus koraiensis*, which have been successfully replanted in secondary forests to aid in SMMFs recovery (Lee et al. 2004; Kiselyova et al. 2021; Wang et al. 2021). To ensure the sustainable development of restored forests, it is crucial to combine restoration techniques with effective protection measures (Kiselyova et al. 2021). Protecting the diversity and integrity of forest habitats is key to maintaining both species diversity and the stability of SMMFs communities (Lin et al. 2024).

## Conclusions

The present study showed that the machine learning approach for classification of mixed coniferous-deciduous forests enables automatic classification with reasonable accuracy, but requires significant sampling and training efforts. Different mixtures of deciduous and coniferous plant populations with different stand densities and life stages may lead to underestimation of vegetation cover using remote sensing. Despite the high mapping accuracy, our results showed large misclassification when SMMFs were detected at intermediate stages of the mapping process. Therefore, ground validation and sampling with visual expert interpretation are needed to improve such automatic classification.

The SMMFs map created in this study, with a high overall accuracy of 0.97 ± 0.03, enables the precise identification of these temperate mixed forest areas across their entire range. This map provides essential insight into the current distribution of SMMFs. There is a noticeable trend of SMMFs being preserved within protected areas throughout their range. Historical anthropogenic activities have led to a significant reduction in the extent and area of SMMFs and the high fragmentation of these forests.

Today, many areas once dominated by SMMFs have been replaced by secondary forests or agricultural lands and anthropogenic non-forest vegetation. For the purpose of SMMFs restoration, it is necessary to design new nature-based solutions for protection, targeted restoration and maintaining biological corridors which would improve the connectivity of forest patches and contribute to the biodiversity conservation in the region. The newly created SMMFs map serves as a valuable tool for these purposes.

## Appendix 1

**Table A1.** The impact of noise on mapping accuracy: overall accuracy (OA), user accuracy (UA), and producer accuracy (PA) ± standard errors (SE) for each country. Optimal results for each country are marked with an asterisk

|  | OA (SE) | UA (SE) | PA (SE) |
| --- | --- | --- | --- |
| **Without filtration** |  |  |  |
| **Russia\*** | 0.98 (0.01) | 0.94 (0.01) | 0.85 (0.1) |
| **China\*** | 0.99 (0.01) | 0.55 (0.05) | 0.51 (0.3) |
| **North Korea** | 0.99 (0.01) | 0.45 (0.04) | 0.40 (0.2) |
| **South Korea** | 0.99 (0.002) | 0.48 (0.05) | 0.99 (0.001) |
| **Filter threshold: 1 pixel** |  |  |  |
| **Russia** | 0.96 (0.02) | 0.95 (0.01) | 0.68 (0.1) |
| **China** | 0.99 (0.01) | 0.57 (0.03) | 0.42 (0.2) |
| **North Korea\*** | 0.98 (0.01) | 0.49 (0.03) | 0.27 (0.2) |
| South Korea | 0.99 (0.002) | 0.49 (0.04) | 0.97 (0.03) |
| **Filter threshold: 2 pixels** |  |  |  |
| Russia | 0.99 (0.01) | 0.95 (0.01) | 0.56 (0.1) |
| China | 0.99 (0.01) | 0.58 (0.03) | 0.09 (0.2) |
| North Korea | 0.98 (0.01) | 0.51 (0.03) | 0.07 (0.2) |
| South Korea* | 0.99 (0.001) | 0.51 (0.04) | 0.99 (0.0001) |
| **Filter threshold: 5 pixels** |  |  |  |
| Russia | 0.99 (0.01) | 0.97 (0.01) | 0.42 (0.1) |
| China | 0.99 (0.01) | 0.61 (0.03) | 0.10 (0.2) |
| North Korea | 0.98 (0.01) | 0.53 (0.03) | 0.03 (0.2) |
| South Korea | 0.98 (0.01) | 0.56 (0.04) | 0.11 (0.05) |

**Table A2.** Mapped area (MA) and estimated area (EA) of SMMF. 95% confidence intervals in km² are given for the estimated area.

| Strata |  | «Loss» | «Gain» | «Stable» | Actual area |
| --- | --- | --- | --- | --- | --- |
| **Total area** | MA | 992.1 | 833.1 | 380.6 | 2178.3 |
|  | EA based on unique sample |  |  |  | **2791.5 ± 1496.2** |
|  | EA based on the sum of areas in individual countries |  |  |  | 3724.8 ± 2293.9 |
| **Russia** | MA | 295.4 | 256.5 | 288.4 | 840.3 |
|  | EA |  |  |  | 1182.4 ± 427.6 |
| **China** | MA | 233.6 | 253.4 | 51.1 | 538.1 |
|  | EA |  |  |  | 746.4 ± 620.1 |
| **North Korea** | MA | 321.7 | 267.3 | 28.1 | 617.1 |
|  | EA |  |  |  | 1691.5 ± 1229.4 |
| **South Korea** | MA | 139.5 | 54.9 | 13.0 | 207.4 |
|  | EA |  |  |  | 104.5 ± 16.8 |

## Supporting Information

Appendix S1. Training dataset

Appendix S2. SMMFs fragments in high-resolution satellite imagery

